# Eutrofication impacted temperate forest vegetation recently

**DOI:** 10.1101/103721

**Authors:** Gabriela Riofrío-Dillon, Jean-Claude Gégout, Romain Bertrand

## Abstract

1. Nitrogen (N) is a key nutrient elements for ecosystems which has been highly impacted by human development and activities since the early 20^th^ century. Despite that changes in N-availability have been demonstrated to impact forests, we still miss evidence of its effect on species composition over the long-term.
2. Based on a large number of floristic observations (n = 45 604), the French forest status related to soil N-availability was reconstructed from herb species assemblage between 1910 and 2010 using both a bioindication approach and a spatiotemporal sampling aiming to pair past and recent floristic observations.
3. We showed that soil N content bioindicated from forest herb communities was higher at the start of the 20^th^ century than over the 2005–2010 period. It decreased more or less continuously until 1975 and 2005 in coniferous (mean ΔC:N=+0.79) and broadleaved (mean ΔC:N=+0.74) forests, respectively, and then was lower than the most recent bioindicated N level observed over the 2005–2010 period (mean ΔC:N=−0.10 and −0.16, respectively). Spatial analysis confirmed the temporal trends with a decrease and increase in forest surface areas where soil N impoverishment and enrichment have been bioindicated over the time, respectively.
4. N bioindicated trends are opposite to changes in N atmospheric deposition compared the 2005–2010 period, while they follow temporal variation in mean N deposition until 1990. *Synthesis*. Our results showed that forest herb communities have been reshuffled in regards of their soil N requirements over the 20^th^ century highlighting that temporal changes in soil N supply have impacted the understory species composition of forest. We evidenced changes in communities towards less nitrophilous plant assemblage followed by a recent eutrophication since 2005. We propose that the nitrogen forest vegetation status is likely related to N atmospheric deposition trend, but also to both acidification, climate change and forestry management which impacted organic matter decomposition and soil N mineralization through effects on soil microbial and fauna activities. The current eutrophication observed in forest herb communities is worrisome for temperate forest ecosystem and its functioning in regards of biodiversity homogenization which often accompanied such a community reshuffling.

## Introduction

Humans activities have transformed the global nitrogen (N) cycle at a record pace (Galloway et al., 2008). Over the last half-century, anthropogenic emissions of N compounds to the atmosphere have overtook emissions from natural processes (Galloway, 2001), and therefore N deposition has increased worldwide (van Aardenne, Dentener, Olivier, Goldewijk & Lelieveld, 2001; Bobbink et al., 2010). Nitrogen is a key element in ecosystems functioning (Vitousek et al., 1997), and its observed increase over time have widely influenced on terrestrial, freshwaters and marine ecosystems (Aber et al., 1998; Carpenter et al., 1998; Schöpp, Posch, Mylona & Johansson, 2003; Clark et al., 2013). The nutrient enrichment caused by N deposition has given rise to changes in soil chemistry (Kristensen, Gundersen, Callesen & Reinds, 2004) which in turn influenced on species yield and ground layer vegetation (Falkengren-Grerup & Eriksson, 1990; Falkengren-Grerup et al., 2000). In forest ecosystems, N increase led to changes in N cycling processes (McNulty, Aber & Boone, 1991; O’Sullivan, Horswill, Phoenix, Lee & Leake, 2011), soil eutrophication and/or acidification (Thimonier, Dupouey, Bost & Becker, 1994; Riofrío-Dillon, Bertrand & Gégout, 2012), tree growth enhancement and higher forest productivity (Solberg et al., 2009; Bontemps, Hervé, Leban & Dhôte, 2011), reshuffling of plant communities (Smart, Robertson, Shield & Van de Poll, 2003), species decline and/or loss (Stevens, Dise, Mountford & Gowing, 2004; De Schrijver et al., 2011), and to increase plant susceptibility to other biotic or abiotic stress factors (Matson, Lohse & Hall, 2002).

After the peak of N deposition in 1980s and the strengthening of implemented policies to control and reduce emissions of nitrogen oxides in 1988, a decrease of approximately 29% et 39% in reduced (NHy) and oxidized nitrogen (Nox) depositions, respectively, was reported for the period 1990–2010 in Europe (EMEP, 2011). In France, N atmospheric deposition followed this trend (EMEP, 2011), but deposition remained at a high level. The effects of the ongoing input of N deposition on ecosystems can still be observed (Bobbink et al., 2010; Thimonier et al., 2010; De Schrijver et al., 2011). It was suggested that not all ecosystems respond to N deposition similarly. Some of them are more susceptible to N excess (Aber, Nadelhoffer, Steudler & Melillo, 1989; Matson, Lohse & Hall, 2002). Elevated N input over time may lead to increase N concentrations in plants and soils, and thus to a release of protons in the soil solution through ammonium (NH_4_^+^) uptake or nitrification (Thimonier et al., 2010). Factors as species composition, forest type, soil, climate, N retention capacity, litter quality, land use history contribute to variation in ecosystem responses to N deposition (Fenn et al., 1998; Matson, Lohse & Hall, 2002; Emmett, 2007). In Europe, there is large variation in nutritional conditions across forest, which is not strongly related to current or historic N deposition (Dise, Matzner & Forsius, 1998). In general, broadleaved forests are more abundant on fertile and N-rich soils, whereas coniferous dominate on less fertile, N-poor soils (Kristensen, Gundersen, Callesen & Reinds, 2004).

Both species response to environmental change (Bobbink et al., 2010) and the limited historical data with measurements of soil parameters highlight the potential usefulness of species composition to bioindicate the environmental conditions and monitoring its changes over the long-term (Braak & Juggins, 1993; Riofrío-Dillon, Bertrand & Gégout, 2012). Among species diversity, plants have demonstrated powerful ability to indicate environmental conditions (Bertrand et al., 2011; Riofrío-Dillon, Bertrand & Gégout, 2012). For instance, significant insight into changes in soil N availability has been shown using floristic data, from forest inventories and/or ancient phytosociological studies, for any time period (e.g. Thimonier, Dupouey & Timbal, 1992; Smart, Robertson, Shield & Van de Poll, 2003). It was suggested that the study of temporal changes in species composition over the shortterm could provide unclear trends of changes (Thimonier et al., 2012), and as a consequence unclear trends of any bioindicated environmental conditions. But such a bioindication approach seems appropriate to detect the effects of N atmospheric deposition or any causes of variation in soil N content on vegetation over the long-term and at large-scale while soil chemistry data is missing (Riofrío-Dillon, Bertrand & Gégout, 2012).

Most of the plant species diversity is represented by the herb layer vegetation in temperate forest ecosystems. It has been shown that herb species have a high sensitiveness to disturbances and therefore a more rapidly and specifically response to them across broad spatial and temporal scales (Falkengren-Grerup et al., 2000; Gilliam, 2007). Then, its dynamics could reflect the evolution of forest status over time (Thimonier, Dupouey & Timbal, 1992; Riofrío-Dillon, 2013). Previous studies have shown significant shifts in plant communities due to direct (Maskell, Smart, Bullock, Thompson & Stevens, 2010; McClean, van den Berg, Ashmore & Preston, 2011) or indirect (through soil mediated effects) effect of N deposition (Stevens et al., 2011a). In France, N atmospheric deposition has deeply influenced changes in environmental conditions over time. These changes have been inspected through vegetation analysis from which a simultaneous forest acidification and eutrophication have been pointed out locally (e.g. Thimonier, Dupouey, Bost & Becker, 1994).

Here, we reconstructed and analyzed the spatiotemporal changes in soil N vegetation status from 45 604 floristic inventories well-distributed across coniferous and broadleaved French forests between 1910 and 2010. Our study was based on forest herb assemblages and their bioindicator character regarding the soil C:N ratio (a common index used in forestry and agricultural sciences to quantify soil N mineralization and soil N content; e.g. Andrianarisoa, Zeller, Dupouey & Dambrine, 2009). It assumes that species follow their nutritional requirements changing their geographical distribution in accordance with changes in nutrient soil conditions to conserve in that way their edaphic niche (Bobbink, Hornung & Roelofs, 1998; Turkington et al., 1998; Riofrío-Dillon, Bertrand & Gégout, 2012). Under this assumption, we expect that N atmospheric deposition have driven plant species reshuffling in forest since the early 20^th^ century, and as a consequence bioindicated N from plant assemblages should follow N deposition spatiotemporal trends. The specific questions we address are: (i) What is the long-term trend in plant communities changes related to soil N in French forests? (ii) Is bioindicated N trend mirroring the N deposition? (iii) Is there a regionalized pattern of bioindicated N? We finally discuss the pattern observed in regards of the N atmospheric deposition but also considering both the increase in litter production, climate warming, changes in soil acidity and forest management since 1910.

## Materials and methods

### Study area

Our study area is the French forest territory which covers 29.2% of Metropolitan France, i.e. a surface of 161 000 km^2^ as determined from the national grid of CORINE Land Cover 2006. French forests are covered by different tree species, which allows the discrimination between pure coniferous and broadleaved forests, and mixed-species forests. Coniferous dominate on less fertile, N-poor soils and are mainly located in mountain range. Broadleaved forests cover more than twice the area of coniferous forests and are mainly located on sites characterized by fertile, N-rich soils (Fig. 1). Because both coniferous and broadleaved forests respond differently to N deposition (Kristensen, Gundersen, Callesen & Reinds, 2004) and herbaceous community can differ substantially (Barbier, Gosselin & Balandier, 2008), we chose to analyze change in plant community related to soil N in these two forest types separately. The forest type information was extracted from CORINE land cover grid 1990 and 2006 for floristic plots sampled before and after 1990, respectively.

**Figure 1:**
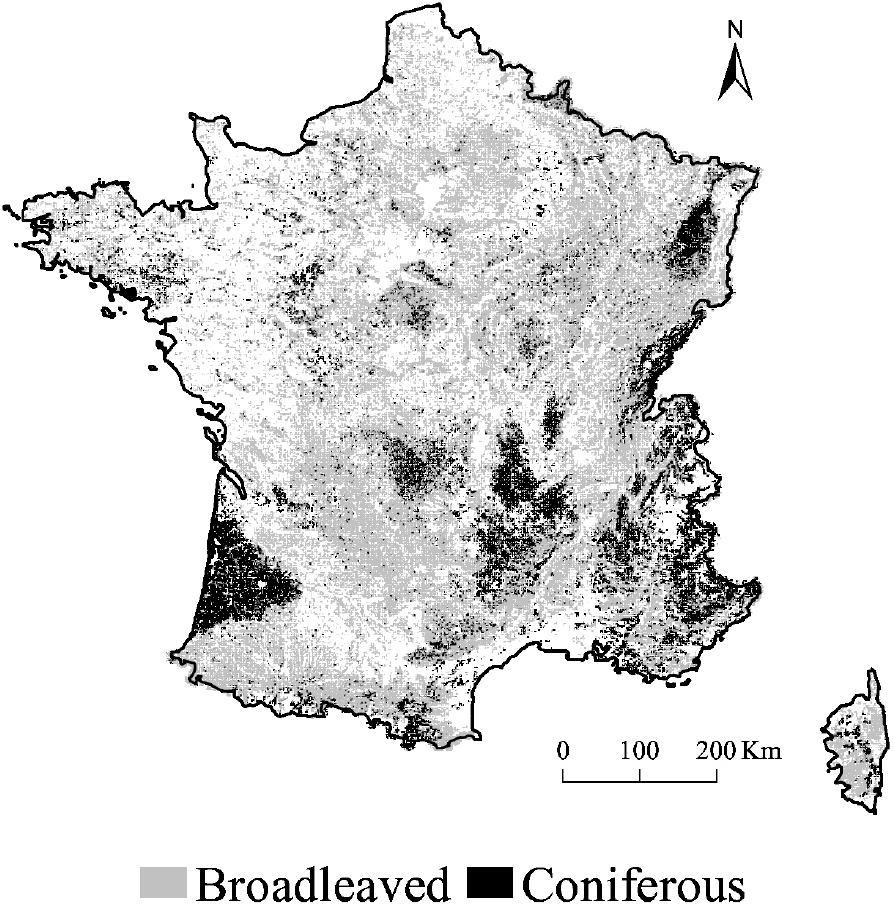
Spatial distribution of coniferous and broadleaved forests throughout Metropolitan France. Coniferous forests cover 22% of forest territory, and broadleaved forests cover 56% of forest territory. Mixed f17,60 cmorests (nor represented) cover the remaining 22% of forest territory. Forest cover is based on CORINE Land Cover 2006.

In France, a low N atmospheric deposition occurred during the first half of the 20^th^ century, followed by a strong anthropogenic deposition since 1950, ranging from 343 mg Nh_y_.m^−2^.yr^−1^ and 108 mg NO_x_.m^−2^.yr^−1^ in 1910 to maximal values of 730 mg NH_y_.m^−2^.yr^−1^ and 763 mg NO_x_.m^−2^.yr^−1^ in 1980s (EMEP, 2011). Then, N deposition have slightly reduced to finish by being constant at a high level.

### Floristic and environmental data

Three databases of floristic inventories with presence/absence data were used: EcoPlant (Gégout, Coudun, Bailly & Jabiol, 2005), Sophy (Brisse, de Ruffray, Grandjouan & Hoff, 1995), and National Forest Inventory (Robert et al., 2010). Together they provided a total of 113 406 floristic plots covering the French coniferous and broadleaved forests, spanning from 1910 to 2010. They included sampling year and location information with an uncertainty less than 1 km.

EcoPlant is a phytoecological database including 4 544 floristic plots between 1910 and 2010; of which 2 483 plots include soil C:N ratio measurements. C and N were measured in the laboratory from the upper organo-mineral A horizon of sampled soils. Total N and organic carbon were determined by Kjeldahl and Anne methods, respectively (Kjeldahl, 1883; Anne, 1945). Sophy is a phytosociological database that includes 24 850 coniferous or broadleaved forest plots from 1915 to 2000. Most sampled plots from these two databases presented an area of 400 m^2^, consistent with current phytosociological practice. The NFI database includes 84 012 floristic plots spanning from 1987 to 2010, which consisted of a surface area of 700 m^2^. Given that each database contains taxonomic issues, a homogenization procedure was carried out to check and update the names of all plant species based on a French nomenclature (TAXREF v5.0; Gargominy et al. 2012). To avoid misidentification issues, we mostly focused on the species level.

### Training, validation and prediction floristic datasets

The selected floristic plots were divided into 2 datasets: the training, comprising 2 483 plots with measured soil C:N ratio and floristic inventories from EcoPlant database, and the prediction dataset, comprising 110 923 available plots only with floristic inventories (Fig. 2a). Floristic plots included in the training dataset were sampled between 1965 and 2010. To minimize the over sampling of some geographic regions and environmental conditions, plots had to be at least 500 m distant from each other. To ensure both a good fit of the N biodindication model (which increases with the species frequency; ter Braak & Juggins, 1993) and the use of a large dataset and pool of species to maximize the spatiotemporal representativeness of our study, species with more than five occurrences were selected resulting in a total of 451 forest herb species for calibration.

**Figure 2:**
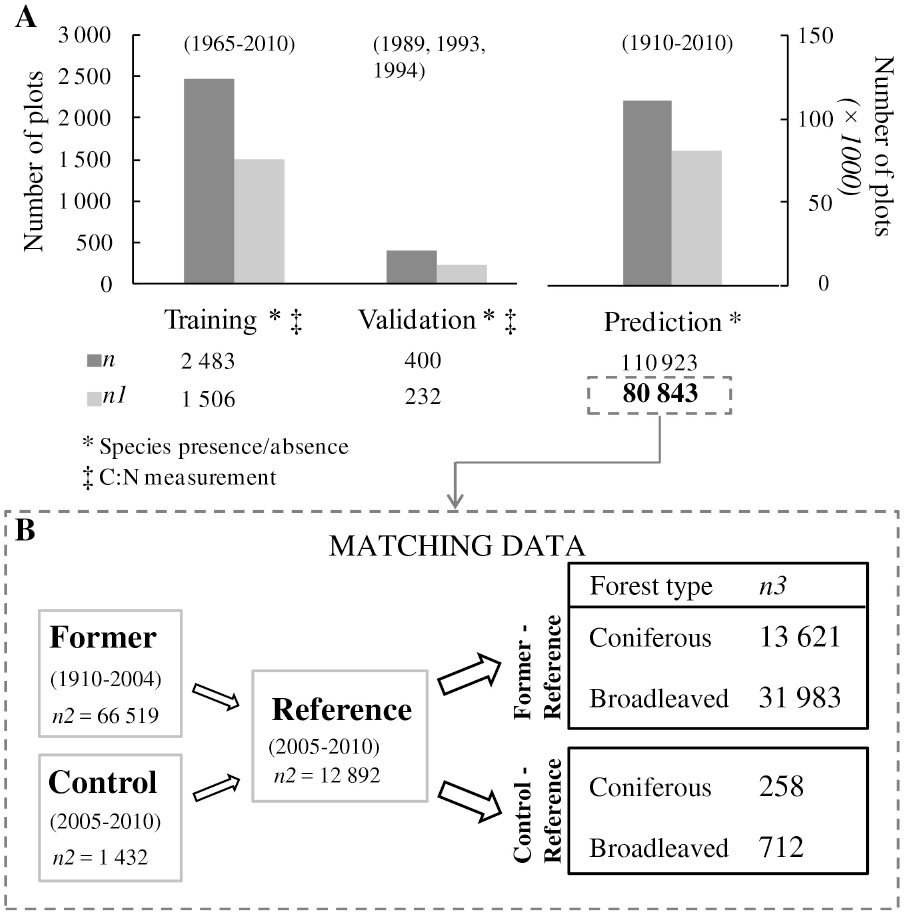
Synthetic representation of the different datasets used. Initial and final number of plots in the training, validation and prediction datasets (a). Scheme of the matching data process in the analysis of temporal bioindicated C:N trends based on the prediction dataset (b). *n* refers to the total initial number of plots, *n1* refers to *n* after filtering plots according to the criteria of selection: species with ≥5 occurrences and ≥3 species per plot, *n2* refers to the number of plots in the prediction sub-datasets, *n3* refers to the number of matched plots analyzed according to the forest type, and after considering the threshold of distance of 3.5 km between matched plots.

Following Riofrío-Dillon et al. (2012), we used a minimum number of three species per plot with at least 5 occurrences into the database for calibrating our predictive model and inferring the C:N value of each floristic plot. These criteria led to a selection of 1506 (i.e. 60.7%) of the initial 2 483 plots of the training dataset that were used to calibrate the C:N bioindication model (Fig. 2a). To assess the performance of the model to infer soil C:N ratio from the forest herb assemblages with an independent dataset, French plots from the 16 x 16 km Network (Badeau et Landmann, 1996) were used as the validation dataset. Bioindicated C:N values were computed from 232 plots (of the initial 400 plots) that met the defined criteria of selection (Fig. 2a). The prediction dataset was used to reconstruct bioindicated C:N trends between 1910 and 2010, To avoid overprediction from our model, plots in the prediction dataset that met the defined criteria of selection were selected, resulting in 80 843 (72.9%) of the initial 110 923 floristic plots (Fig. 2a).

### Weighted Averaging Partial Least Squares (WA-PLS) regression to infer soil N availability from floristic assemblages

Among several available techniques using biotic data as a tool for reconstructing past environmental variables, WA-PLS is a powerful inverse approach, i.e. that the adjusted model predicts directly environmental variables as a transfer function from species assemblages with some error (ter Braak & Juggins, 1993). WA-PLS is appropriated for calibration when the species-environment relationship are unimodal (i.e. one optimum in the ecological niche space), and the species data are binary (presence/absence) (ter Braak, 1995). WA-PLS is a combination of both weighted averaging (ter Braak & Barendregt, 1986) and PLS methods (ter Braak, 1995). It can be summarized as follows:

i. The training dataset is first transformed to linearize species-environment relationship: species assemblage dataset is weighted by both the number of species per plot and the species frequencies (*x*\*), and the environmental variable is weighted by the number of species per plot (*y*\*).
ii. A PLS regression is conducted on the transformed training dataset to fit linear combinations (*f()* or principal components of the PLS) of the predictors (*x*\*) so as to maximize the prediction of the environmental variable: *y*\* = *f*(*x*\*) + *error*. PLS regression produces an initial component as a set of coefficients, or weighted averages of the species optima with respect to a given environmental variable. The second and further components are selected by optimizing the prediction of the environmental variable (*y*\*) as for the first component, and use the residual structure in the data to improve the estimates of the species optima (each new component is orthogonal to the previous one) (ter Braak, 1995). The number of components that gives the best transfer function requires an examination of performance statistics generated by leave-one-out cross-validation (Birks, 1998), and a confirmation on an independent dataset (ter Braak, 1995).
iii. Postprocessing transformation of the results of the PLS regression is required to predict values of the environmental variables (for more details read ter Braak & Juggins, 1993). WA-PLS is a training procedure that has already been successfully used in pollen and species assemblages analyses to reconstruct past climatic (Bertrand et al., 2011) and edaphic conditions (Riofrío-Dillon, Bertrand & Gégout, 2012).

Here, the WA-PLS approach was used to infer soil C:N ratio from herb assemblages. A WA- PLS model was calibrated linking the floristic assemblage (among a pool of 451 species) of each of the 1 506 plots in the training dataset with their corresponding measured soil C:N ratio. A 5-component WA-PLS model was selected on the basis of its root-mean-square deviation (RMSD = 2.42), low bias (mean of prediction error [bioindicated – measured C:N] = −0.009 C:N units) and coefficient of determination between observed and predicted values on the validation dataset (R^2^ = 0.574; Fig. 3). Using the prediction dataset, the bioindicated C:N values were inferred from the species assemblages for each floristic plot based on the calibrated 5-component WA-PLS model.

**Figure 3:**
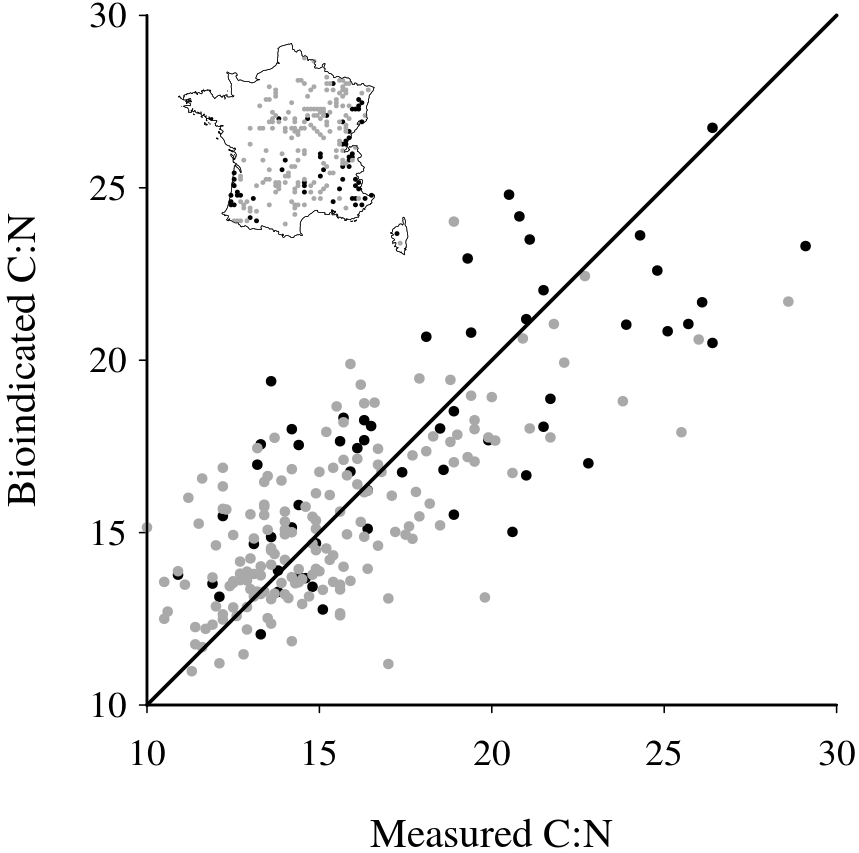
Relationship between measured soil C:N values and bioindicated C:N values (predicted from the 5-component WA-PLS model) from the validation dataset (R^2^ = 0.574; mean difference = −0.009 C:N units; RMSD = 2.42; *n* = 232 surveys). Black and gray points represent the spatial distribution of surveys and C:N values in coniferous and broadleaved forests, respectively. Solid black line represents *y* = *x*.

### Temporal sampling for analysis of forest vegetation status in regards of soil nitrogen changes

To determine changes in the soil N availability over time, the prediction dataset was divided into five periods considering the significance of air pollution between 1910 and 2010 as well as the data availability. The first period (1910–1949, *n* = 1 043 plots) was defined as the earliest period, with a low deposition of approximately 379 mg Nh_y_.m^−2^.yr^−1^ and 179 mg NO_x_.m^−2^.yr^−1^ in average across the French territory (EMEP, 2011). The second period (1950–1974, *n* = 5 186 plots) has been described as a period of increasing N pollution (on average 521 mg NH_y_.m^−2^.yr^−1^ and 458 mg NO_x_.m^−2^.yr^−1^ deposited; EMEP, 2011). The third period (1975–1989, *n* = 14 324 plots) was defined as a period of the highest N atmospheric deposition. Moreover, at that time environmental policies were implemented to control air pollution (i.e., Convention on Long-range Transboundary Air Pollution, Geneva, 1979). The fourth period (1990–2004, *n* = 45 966 plots) was defined as a period of slight reduction of N deposition. The fifth period (2005–2010, *n* = 14 324 plots) was defined as a period of no change in N deposition, remaining however high.

To reconstruct long-term spatiotemporal changes of forest vegetation status related to soil N in the absence of permanent surveys, a method that allows the comparison of floristic plots over time was used (cf. Riofrío-Dillon, Bertrand & Gégout, 2012). The floristic plots collected most recently (i.e. between 2005 and 2010) were defined as the “reference” data which represents a large number of available floristic plots well-distributed across French forest territory. The 66 519 remaining floristic plots spanning from 1910 to 2004 were defined as “former” data and used to compute bioindicated C:N changes regarding the “reference” plots. To control for the potential effect of spatial variability on our assessment of soil bioindicated C:N changes between periods (as described below), 10% of “reference” data were extracted randomly to provide “control” plots (n = 1 432 plots). Consequently, 12 892 “reference” plots were used in the matching process (Fig. 2b).

The method used consisted of matching each plot from the 1910–1949, 1950–1974, 1975–1989, and 1990–2004 periods (i.e. “former” data) with the nearest plot from the “reference” data, with both plots located on the same forest type. The nearest neighbor was determined by computation of the Euclidean distance (d) between floristic plots. The bioindicated C:N temporal change was computed for each pair (Δ*C:N* = *C:N_reference_* – *C:N_former_*) and separate analyses were conducted for both coniferous and broadleaved forests.

Because we aimed to minimize the ΔC:N between matched plots due to geographical distance, the “reference” plots were used to explore the spatial autocorrelation between bioindicated C:N values. Then, a threshold of distance was defined to select plots sufficiently closed to each other to allow robust temporal analyses. The spatial autocorrelation between bioindicated C:N values with respect to the forest type was analyzed using variograms (Fig. S1) (Fortin & Dale, 2005). Considering the variogram outputs, a threshold of distance less than 3.5 km between “former” and “reference” plots was selected because bioindicated C:N values within this radius were spatially autocorrelated. Considering this comparative distance radius, a total of 45 604 matched plots were obtained, of which 13 621 were situated in coniferous forests (median d_coniferous_ [1st to 3rd quartile] = 1.8 [1.1–2.5] km) and 31 983 in broadleaved stands (median d_broadleaved_ [1st to 3rd quartile] = 1.8 [1.0–2.4] km) (Fig. 2b). Such distance criteria between matched floristic observations limit changes in bioindicated C:N values due to spatial variation in sampling which could bias our temporal trends. The statistical significance of the ΔC:N between periods (i.e. ΔC:N_t_ vs. ΔC:N_t+1_; with t defining a period) and the statistical significance of the ΔC:N of a period per se (ΔC:N_t_ vs. ΔC:N_t_ = 0) were both tested using Wilcoxon Rank Sum test (*P* < 0.05).

To assess the validity of our method, distinguishing temporal and geographical variations, the “control” plots were matched to the nearest “reference” plot (as described above). Considering the threshold of distance less than 3.5 km between “control” and “reference” plots, a total of 970 matched plots were obtained, of which 258 were located in coniferous forests and 712 in broadleaved forests (Fig. 2b). As each pair consisted of two floristic plots belonging to the reference period, no differences of bioindicated C:N within “control”-”reference” pairs were expected in the absence of sampling bias in our matching method. According with this expectation, the ΔC:N did not significantly differ from 0, in either coniferous (mean ΔC:N = 0.13 C:N units [SE = 0.22], *P* = 0.446) or broadleaved forests (mean ΔC:N = 0.02 C:N units [SE = 0.11], *P* = 0.727). Further, the median distances between “control”-“reference” pairs was 2.0 km in both coniferous and broadleaved forests, which is comparable with distances reported between “former” and “reference” matched plots. Hence, the suitability of our method was validated, indicating no spatial bias.

### Sampling for analysis of the spatial variation of N availability in forest soils

To visualize where bioindicated C:N temporal changes had occurred across the French forest territory, we performed a spatial reconstruction. The floristic plots of the prediction dataset were mapped, differentiating both coniferous and broadleaved forests and considering 1910–1949, 1950–1974, 1975–1989, and 1990–2004 periods. Spatial reconstructions were based on the 50 × 50 km EMEP grid (EMEP, 2011). A total of 319 grid cells cover Metropolitan France. For each grid cell and period, mean bioindicated ΔC:N values were computed based on at least five matched plots between the former and reference periods (see above for details about the matching data method used). The statistical significance of the bioindicated C:N changes by cell (C:N_i,t_ vs. C:N_i,ref_ with i defining a cell with bioindicated C:N values in both compared periods, and t defining a former period) was tested using Wilcoxon Rank Sum test (*p* < 0.05). Separate analyses were conducted for coniferous and broadleaved forests.

### Assessment of the effect of nitrogen atmospheric deposition on bioindicated C:N temporal changes

Nitrogen atmospheric deposition (nitrogen oxyde, ammonia and total nitrogen) has been predicted at the location of every matching floristic observations from the EMEP model (EMEP, 2011). Temporal changes in N atmospheric depositions were computed for each pair as the difference between N depositions during the reference and former periods (ΔNdep = Ndep_reference_ – Ndep_former_; as for ΔC:N). Mean N atmospheric deposition between the reference and former periods was also computed for each pair of floristic observations. Temporal trends of these two variables were finally compared to biondicated C:N temporal changes in coniferous and broadleaved forests, and their effects were tested using a linear model (Student’s *t* test comparing the effects of both temporal changes in N deposition and mean N deposition to 0).

All models and statistical analyses were performed in the *R* environment (R Development Core Team, 2013). We used the “pls” package (Mevik & Wehrens, 2007) and personal codes to calibrate and predict bioindicated C:N values. We used ArcGIS and its Geostatistical Analyst extension for spatial and geostatistical analysis (version 10.1; ESRI Inc., Redlands, CA, USA).

## Results

### Analysis of temporal bioindicated soil C:N trends

The reconstructed bioindicated trends of ΔC:N showed similar magnitudes of changes between coniferous and broadleaved forests over time (Fig. 4). In coniferous forests, the bioindicated C:N changes between past and the 2005–2010 periods increased significantly by +0.75 C:N units in average (SE = 0.12; *P* < 0.01) between 1910 and 1990, with a high increase from 1950 (mean ΔC:N = +0.38 C:N units [SE = 0.08], *P* < 0.001; Fig. 4a). During this phase, bioindicated C:N changes switched from a period of N impoverishment (highlighted by positive ΔC:N values in Fig. 4a) to a period of N enrichment from 1975 (highlighted by negative ΔC:N values in Fig. 4a). After 1990, bioindicated C:N changes stabilized at a level of N enrichment close to the one reported between 1975 and 1990. Then, the bioindicated C:N changes evidenced an eutrophication of forest plant communities in the most recent period studied, i.e. a decrease in bioindicated C:N values since 2005 (mean ΔC:N = −0.23 C:N units [SE = 0.03], *P* < 0.01; Fig. 4a).

**Figure 4:**
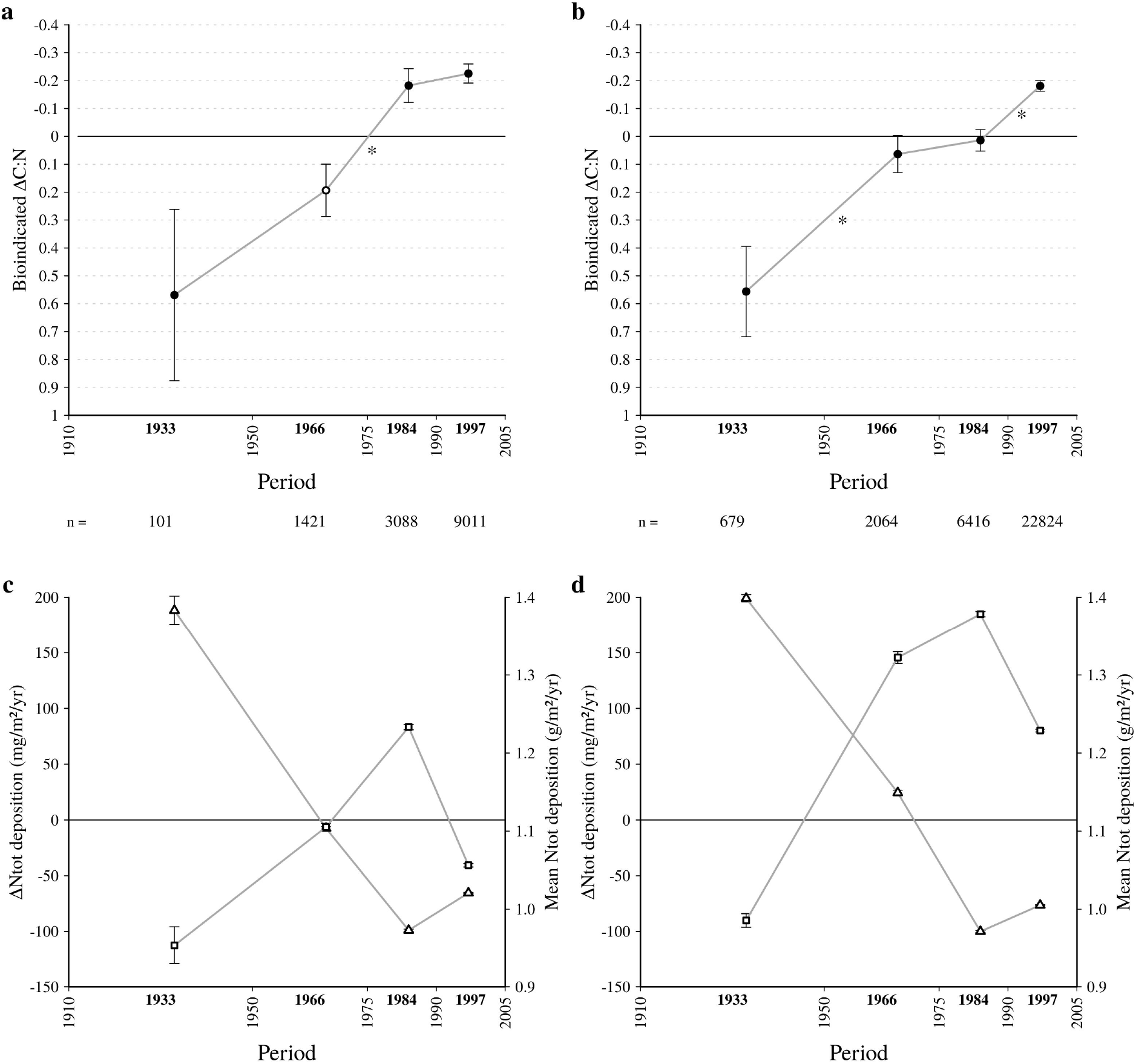
Temporal bioindicated soil C:N and N atmospheric deposition changes between 1910 and 2005 in (a, c) coniferous and (b, d) broadleaved forests. ΔC:N has been computed as the bioindicated C:N differences between matched observations of the reference and former periods (ΔC:N = C:N_reference_ – C:N_former_). Two temporal trends of N atmospheric deposition based on predictions of the EMEP model (EMEP, 2011) are shown in panels c and d. Temporal changes in total N deposition has been computed as the difference in depositions between matched observations of the reference and former periods (ΔNtot = Ntot_reference_ – Ntot_former_). Mean total N deposition between the reference and former periods has also been computed. The ΔC:N axis in panels a and b have been reversed in order to depict changes in N and to allow direct comparison with N deposition trends. Values above and below the line of no change (Δ = 0) show higher and lower bioindicated soil N content and N deposition, respectively, during the most reference period studied. Mean values of the bioindicated C:N and N deposition changes, as well as mean N deposition are shown (circle, triangle and square, respectively) with standard error (error bars). The statistical significance of the bioindicated C:N change of a period *per se* is displayed by closed circles (*P* < 0.05; Wilcoxon Rank Sum test comparing ΔC:N values to 0). The statistical significance of the bioindicated C:N changes between periods is displayed by asterisks (*P* < 0.05; Wilcoxon Rank Sum test comparing ΔC:N values of successive periods). Bold dates above the *x*- axis are the mean year of each defined period. The number of matched records (n) analyzed in each period for coniferous and broadleaved forests is displayed below.

In broadleaved forests, the bioindicated C:N has increased significantly by +0.74 C:N units in average (SE = 0.07; *P* < 0.001) between 1910 and 2005 (Fig. 4b). During this period, the trend in bioindicated C:N changes (comparing bioindicated C:N values between past and the 2005–2010 periods) was not continuous. After a first high bioindicated C:N increase between 1910 and 1975 (mean ΔC:N = +0.49 C:N units [SE = 0.11], *P* < 0.001), the trend was null until 1990 and then restarted to increase (Fig. 4b). This last phase was also characterized by a period of N enrichment compared to the reference period (ranging from 2005 to 2010; mean ΔC:N = −0.18 C:N units [SE = 0.02], *P* < 0.001), while N impoverishment was bioindicated by forest plant communities before 1990 (highlighted by positive ΔC:N values in Fig. 4b).

The comparison of bioindicated ΔC:N trends (Fig. 4ab) to changes in N atmospheric deposition (Fig. 4cd) in coniferous and broadleaved forest showed opposite pattern. Bioindicated ΔC:N trends followed the increase in mean N atmospheric deposition trends until 1990 whatever the forest type (Fig. 4). During this period, bioindicated ΔC:N trends have even varied in a same order of magnitude as mean N atmospheric deposition (i.e. both trends have reduced or accelerated similarly). Bioindicated ΔC:N trends showed an eutrophication of floristic assemblage since 2005 while mean N atmospheric deposition decreased at a level lowest than the one recorded between the 1950–1975 and 2005–2010 periods.

### Analysis of spatial variation in temporal bioindicated soil C:N changes

The spatial analysis showed that a largest cover of the French forests did not experiment any changes in bioindicated C:N between before and after 2005 (Fig. 5). However, at least one significant bioindicated C:N change has been observed in 99.2 and 100% of the broadleaved and coniferous forest areas studied, respectively, between 1910 and 2010. In agreement with the global trends reported in Fig. 4ab, the spatial analysis showed a transition from a period of N impoverishment to a period of N enrichment (Fig. 5). This eutrophication of the forest communities happened early in the coniferous forests as the surface area where bioindicated C:N decreased began to be larger than the area where bioindicated C:N increased from 1950. In the deciduous forests, this transition was observed from 1990.

**Figure 5:**
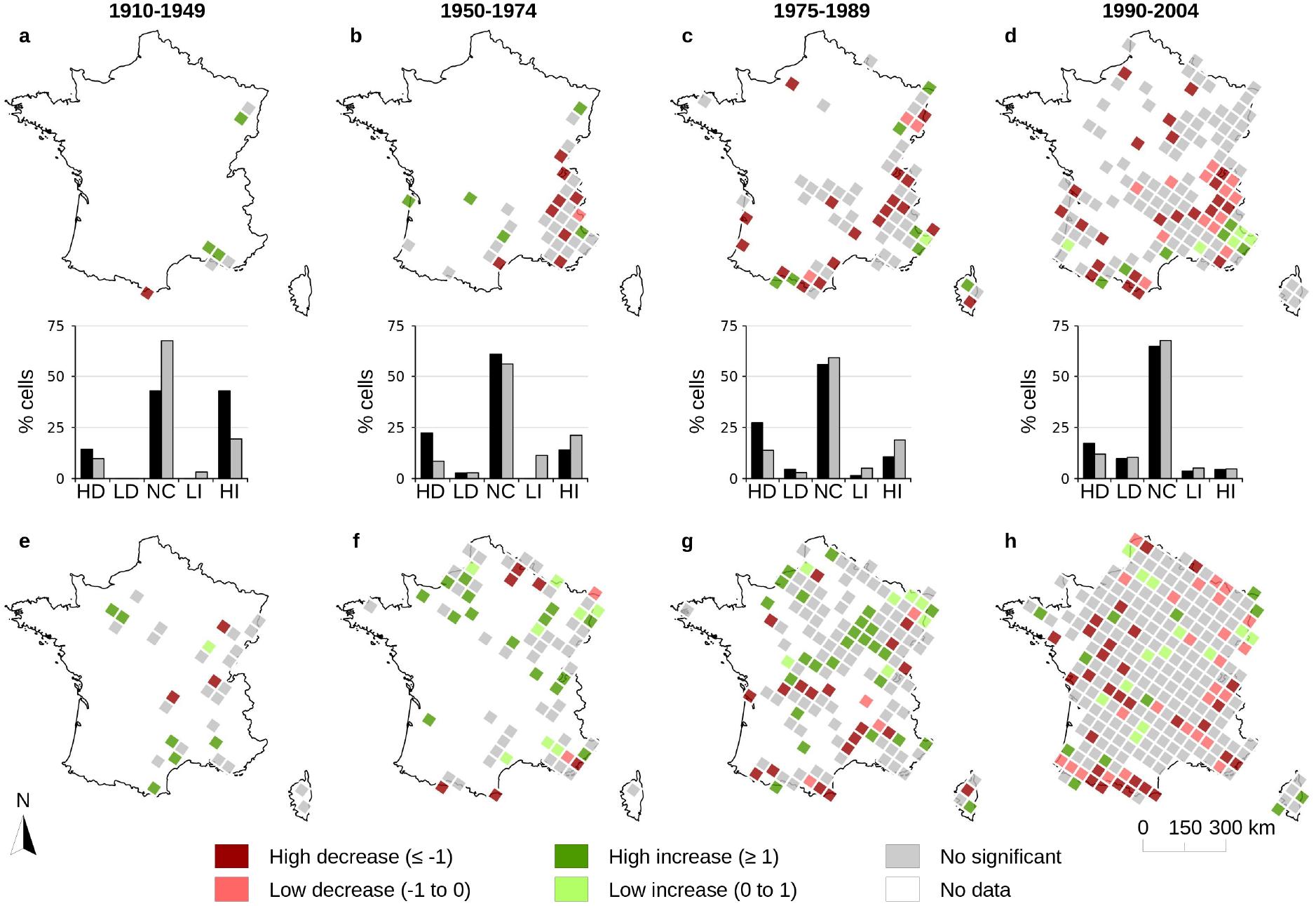
Spatiotemporal bioindicated soil C:N changes in (a, b, c, d) coniferous and (e, f, g, h) broadleaved forests. Mean values of bioindicated C:N changes per cell between former (time periods are displayed in the up of the figure) and reference periods are shown (ΔC:N = C:N_reference_ – C:N_former_). The significance of the bioindicated C:N changes (*P* < 0.05; Wilcoxon Rank Sum test) is displayed by graduated shades of red for decreasing bioindicated C:N, graduated shades of green for increasing bioindicated C:N and gray color for non significant changes. White indicates no data (i.e. EMEP grid containing less than 5 bioindicated C:N estimates in each period compared). The histograms represent the percent of grid cells for coniferous (black bars) and broadleaved forests (gray bars) showing a high decrease (HD), low decrease (LD), no change (NC), low increase (LI), and high increase (HI) of bioindicated C:N.

The temporal changes in bioindicated C:N varied locally and between each period throughout the French territory whatever the forest type. However, the bioindicated C:N increase in regards of the recent C:N levels (i.e. during the reference period) was mainly observed throughout the northern half of the French broadleaved forest territory between 1950 and 1990. On the contrary, a decrease in bioindicated C:N spread throughout the whole forest territory between the 1990–2004 and 2005–2010 periods. In particular, mountain areas such as Pyrenees (south-western of France) and Alps (southeastern of France) exhibited this pattern. In coniferous forests, the decrease in bioindicated C:N has even started in 1950 in mountain areas (mainly Alps).

Bioindicated C:N changes were mainly not correlated to changes in N atmospheric deposition between before and after 2005 whatever both the forest type and the form of the N deposition (nitrogen oxide or ammonia) considered (Fig. S2 and S3). Changes in bioindicated soil N content (derived from bioindicated C:N) increased with changes in N atmospheric deposition in coniferous forests for the 1910–1949 and 1975–1989 periods, while an opposite relationship was observed in broadleaved forest for the 1910–1949 period (Fig. S3). Changes in bioindicated soil N content were negatively correlated to mean nitrogen oxyde deposition in broadleaved forests (mainly since between 1975; Fig. S4 and S5). In coniferous forests, changes in bioindicated soil N content were positively correlated to mean ammonia deposition (mainly since 1990; Fig. S4 and S5).

## Discussion

Nitrogen is a key nutrient elements for ecosystems which has been highly impacted by human development and activities since the early 20^th^ century (Vitousek et al., 1997; Clark et al., 2013). Despite that changes in soil N supply have impacted forests in different ways, we still miss spatiotemporal evidence of its effect on species composition over the long-term and large spatial scale (mainly because soil chemistry monitoring before 1980 is rare; Bobbink et al., 2010). Here, we reconstructed the spatiotemporal changes in soil N vegetation status from 45 604 floristic inventories well-distributed across the French metropolitan forest territory between 1910 and 2010. Our study was based on forest herb assemblages and their bioindicator character regarding the soil C:N ratio. It assumes that plants track their ecological niche over space and time shifting their geographical distribution and reshuffling the composition of species communities in accordance with changes in soil nutrient conditions (Bobbink, Hornung & Roelofs, 1998; Turkington et al., 1998; Riofrío-Dillon, Bertrand & Gégout, 2012). Given both the importance of the edaphic dimension in the definition of the ecological niche of plants (Bertrand, Gégout and Bontemps, 2011; Riofrío-Dillon, Bertrand & Gégout, 2012; Dubuis et al., 2013) and the high responsiveness of plants to nutrient-richer conditions (Bobbink, Hornung & Roelofs, 1998; Turkington et al., 1998), we argue that our bioindication approach is pertinent, especially under a context of high temporal variation in N input (such as N atmospheric deposition for instance). Indeed, forest plant communities has been demonstrated to be reshuffled rapidly in response to soil eutrophication (Thimonier, Dupouey, Bost & Becker, 1994) and acidification (Riofrío-Dillon, Bertrand & Gégout, 2012). Moreover, predictions of our soil C:N bioindication model were comfortably correlated to soil C:N observations (R^2^=0.574) and their errors (RMSD=2.42) were unlikely sufficient to challenge our results and conclusions based on more than 45 000 floristic observations. The bioindicated C:N trends reported here have also unlikely mirrored other concurrent global changes which could drive plant community reshuffling over the time (such as climate change for instance; Bertrand et al., 2011). It has been recently shown that the signal of concurrent global changes impacting plant communities is present in residuals of the bioindication model (Bertrand et al., 2016). It means that the bioindication model (as the one we used here) is able to disentangle the numerous environmental change dimensions which contribute to explain plant assemblage and their spatiotemporal changes, in order to capture and predict, respectively, the signal and values of the specific environmental dimension studied.

### long-term changes in forest herb communities in response of soil nitrogen temporal changes

Our analyses have shown that the composition of the temperate forest herb communities have changed in regards of their soil N requirement since the early 20^th^ century. Such a long-term reshuffling demonstrates that forest plant communities have responded to change in soil N contents. We reported that soil N content bioindicated from forest herb communities was higher in the early 20^th^ century than over the most recent period (ranging from 2005 to 2010), meaning that former plant communities were more adapted to high soil N content than today. The forest herb assemblages were reshuffled more or less continuously towards species assemblages adapted to poorer soil N content until 1975 and 2005 in coniferous and broadleaved forests, respectively. Then, soil N requirements of plant communities stabilized in coniferous forests, before to increase after 2005 whatever the forest type considered (mean ΔC:N=−0.10 and −0.16 in coniferous and broadleaved forest, respectively). Such a last phase highlights a recent soil eutrophication which has led forest herb communities to reshuffle towards more nitrophilous species assemblages. Despite the present study has no analog studies in term of spatial and temporal scales covered, the recent eutrophication highlighted here by forest plant communities is supported by similar observations made locally in forests (Thimonier, Dupouey, Bost & Becker, 1994; Diekmann, Brunet, Rühling, & Falkengren-Grerup, 1999). However, these previous studies considered this eutrophication phase has started since 1970s in temperate forests which is 35 years earlier than what we observed at the scale of the French forest territory (a general result which varies locally with the forest type as Fig. 5 showed). Such a pattern in plant assemblage has been also demonstrated to be associated in biodiversity loss (e.g. Stevens, Dise, Mountford & Gowing, 2004) and homogeneization (especially in the most N constrained soils, e.g. Reinecke, Klemm & Heinken, 2013; Naaf and Kolk, 2016). It may have important consequences for the ecosystem functioning especially if the functions of extinct species are not sustained by plant species composing the new communities (e.g. Isbell et al. 2011). Finally, our results provide a new evidence that forest plant communities have been highly impacted by global changes over the long-term, and complete recent demonstrations that the composition of these communities have been changed by climate warming (Bertrand et al. 2011) and soil acidification (Riofrío-Dillon, Bertrand & Gégout, 2012) over the 20^th^ century.

### A high level of nitrogen atmospheric deposition as a potential driver

As a potential explaining factor of the bioindicated N trends observed in French forests, we analyzed the correlations with N atmospheric deposition. We showed that changes in N atmospheric deposition between past and the 2005–2010 periods depicted an opposite temporal trends than the one observed for bioindicated N changes. This unexpected relationship means that when N deposition increased between two dates, the soil N content bioindicated from floristic assemblages decreased during the same time period. This observation contrasts with the numerous studies which have highlighted both a loss of plant species and a reshuffling of forest plant communities towards nitrophilous species assemblages in response of the N atmospheric deposition increase over the 20^th^ century (e.g. Thimonier, Dupouey, Bost & Becker, 1994; Stevens, Dise, Mountford & Gowing, 2004; Bobbink et al., 2010; Reinecke, Klemm & Heinken, 2013 ; Dirnböck et al. 2014; Gilliam et al. 2016). However, Both trends in bioindicated N and mean N atmospheric deposition varied synchronously demonstrating that a decrease or increase in N deposition level may induce a similar change in bioindicated N. It means that rather than changes in N atmospheric deposition, it is the mean level of N atmospheric deposition which may explain the temporal trends in bioindicated N. This result is supported by previous observations (e.g. Thimonier, Dupouey, Bost & Becker, 1994; Diekmann, Brunet, Rühling, & Falkengren-Grerup, 1999; Bobbink et al., 2010; Dirnböck et al., 2014), but in the present study the apparent temporal correlations between N deposition and bioindicated N showed a slowdown in the reshuffling of plant communities towards species assemblage adapted to poorer nutrient soil rather than an eutrophication of plant communities. Since 1990, the trends were decoupled with a decrease in mean N atmospheric deposition while the bioindicated N remained stable and even increased since 2005. It demonstrates that floristic assemblages are still responding to the N atmospheric deposition likely due to high level of N input accumulated in forest soils during the past decades. Indeed, N atmospheric deposition, after having reach its peak in 1980s, have stabilized and decreased slowly since 1995 but is stayed high with a deposition level exceeding 10–15 kg.ha”^1^.yr”^1^ considered as the critical loads of temperate forests (Bobbink et al 2010). The repetition of years with N atmospheric deposition close or exceeding critical loads has likely contributed to initiate the recent reshuffling towards more nitrophilous plants assemblages. Furthermore, soils buffer N pollution particularly when both C:N ratio is high and there are large stocks of labile C (Emmett et al., 2010) as we observed since 1975 in French forests. Such a buffering effect has likely contributed to delay the impact of N atmospheric deposition on forest herb communities, and as a consequence to delay its reshuffling towards nitrophilous species assemblage. Finally, our results are also coherent with the spatial distribution of N atmospheric deposition throughout the French territory. In particular, we highlighted forest herb communities over mountain areas such as Pyrenees (south-western of France) and Alps (south-eastern of France) have highly and recently changed towards nitrophilous species assemblages while these areas have been particularly exposed to high nitrogen oxyde and ammonia atmospheric deposition levels since the late 1990s (Croisé, Ulrich, Duplat & Jaquet, 2005).

### Alternative drivers of the bioindicated soil nitrogen content trends

In the present study, we have not directly reconstructed soil N content from forest herb assemblages but rather bioindicated the soil C:N ratio. It means that our results are on the combined dependence of spatiotemporal changes in soil N and organic carbon contents. We assessed the effects of these compounds in our data, and demonstrated that soil N content contributed more to decrease soil C:N ratio (size effect (a) = −1.82; assessed in the following linear model 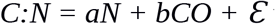 with *a* and *b* are the coefficients of the soil nitrogen (*N*) and organic carbon (*CO*) contents, respectively) than the carbon content is able to increase (size effect (b) = 1.75). It indicates that the soil C:N ratio is more sensitive to variations in soil N content than in soil organic content and confirms our choice to interpret the soil C:N ratio mainly as a proxy of soil N supply. We also consider that the use of bioindicated C:N is advantageous in our study. The soil C:N ratio is thought to be an indicator of the amount of inorganic N in soils which is available for plant uptake and leaching (Gundersen, Callesen & de Vries, 1998; Nave, Vance, Swanston & Curtis, 2009), a key diagnostic factor of ecosystem response to N addition (Adams et al., 2004), a variable that provides information on the relative change in C and N storage in soils (Nave, Vance, Swanston & Curtis, 2009; Emmett et al., 2010) and an important parameter in soil chemistry models (Emmett, 2007). Here, it notably allows us to go beyond the observed changes in soil N supply bioindicated from forest herb assemblages by trying to explain our results considering underlying processes such as organic matter decomposition and soil N mineralization.

The biomass productivity of temperate forests has increased over the 20^th^ century under the impetus of climate change (e.g. Nemani et al., 2003; Boisvenue and Running, 2006) and N atmospheric deposition (e.g. Bontemps, Hervé, Leban & Dhôte, 2011) which have released climate and nutrient limitations on the growth of numerous tree species (e.g. Charru, Seynave, Hervé, Bertrand & Bontemps, 2017). In France, forest management has also contributed to increase forest biomass by favoring forest growing stock at the expense of intensive wood harvesting (IGN 2016; Roux et al., 2017). The increase in forest biomass production implies an increase in litter and thus in soil organic matter. Its accumulation has likely increased the soil organic carbon content over the 20^th^ century. Such a scenario is congruent with the increase in soil C:N ratio evidenced by plant communities until 1990 and 2005 in coniferous and broadleaved forests, respectively. However, organic matter is also a source of N. It means that some factors have constrained soil N mineralization in forests during this period.

Soil acidity and climate conditions are known to impact soil nutrient supply. Soil acidity and acidification inhibit the activities of the fauna and microbial communities in charge of organic matter decomposition and N mineralization in soils (Blagodatskaya & Anderson, 1998; Falkengren-Grerup, Brunte & Diekmann, 1998; Blanco, Wei, Jiang, Jie & Xin, 2012; Liu et al., 2017). Such a process has direct impact on the soil C:N ratio which is evidenced in our data by a negative relationship between the soil C:N ratio and pH in the more acidic soils (Fig. S6). It indicates the important effect of soil acidity on the soil C:N ratio, and also on the underlying processes defining it such as mineralization and decomposition activities of soil fauna and microbial communities. In contrary, climate warming has been reported to mainly increase soil organic matter decomposition (Kirshbaum, 1995; Qualls, 2016) and N mineralization (Wang et al., 2016; Liu et al., 2017) with a greater impact in areas where cold temperatures were limiting soil fauna and microbial activities and where soil moisture has not decreased. In France, acidification of forest soils have been reported from 1910 to 1990 (Riofrío-Dillon, Bertrand & Gégout, 2012) and climate has warmed over the 20^th^ century (especially after 1986; Bertrand et al. 2011). Thus, both soil acidification and coolest temperature conditions experienced by fauna and microbial communities before 1990 have likely reduced litter decomposition and soil N mineralization, and as a consequence led to both litter accumulation and less N released to the soil solution. Such a scenario is congruent with the increase of bioindicated C:N that we observed until 1990. Furthermore, soil acidification has slowed down over the 20^th^ century. Its blocking effect on organic matter decomposition and soil N mineralization had likely been progressively released and may explain the stabilization of the bioindicated C:N started from 1975 as well as the recent eutrophication evidenced by forest herb communities. Such a process has likely been boosted by the high temperature increase since 1986.

### Conclusion

The present study evidences long-term spatiotemporal forest herb communities changes in regards of soil nutrient content. Plant communities bioindicated a decrease in soil N content during the 20^th^ century followed by a recent eutrophication since 2005. N atmospheric deposition was unlikely to explain alone such a trend which seems to be more the result of a set of interacting global change factors. The soil N impoverishment bioindicated by floristic assemblages until 1990 is likely explained by the increase in soil organic matter as a result of forest management and climate change which have boosted forest production and the inhibiting effect of soil acidification on microbial and fauna communities in charge of soil organic matter decomposition and N mineralization. Such a change in plant communities have been progressively mitigated over the 20^th^ century and recently inversed likely through both the accumulation of high N atmospheric deposition in soils and the release of soil acidity and climate constraints on organic matter decomposition and N mineralization activities of soil microbial and fauna communities. The recent eutrophication observed in plant communities is worrisome for temperate forest ecosystem functioning and conservation in regards of biodiversity homogenization which often accompanied such a community reshuffling. It could have important implications in current environmental policies and current permissible levels of N pollutions.

## Acknowledgements

We thank E. Dambrine and D. Allard for comments on a previous version of the manuscript; I. Seynave, H. Brisse, P. de Ruffray and J. M. Frémont for contributions to the EcoPlant, Sophy and NFI databases; and all who participated in the conceptions of these databases. The phytoecological database EcoPlant was funded by the National Institute of Rural, Water and Forestry Engineering (ENGREF, AgroParisTech), the National Forest Department (ONF), and the French Environment and Energy Management Agency (ADEME). The R.B.’s work is supported by the TULIP Laboratory of Excellence (ANR-10-LABX-41). This study was funded through a Ph.D. grant to G.R.-D. by ADEME and Lorraine Regional Council.

## Author contributions

G.R.-D., J.-C.G. and R.B. conceived and designed the study as well as the analytical framework. R.B. and G.R.-D. contributed equally to format the floristic database. G.R.-D. analysed the data with the support and supervision of R.B. G.R.-D. wrote the first draft of the paper. R.B. rewrote later versions of the manuscript. All authors discussed the results and contributed to manuscript revisions.

## Supplementary Information

**Figure S1:**
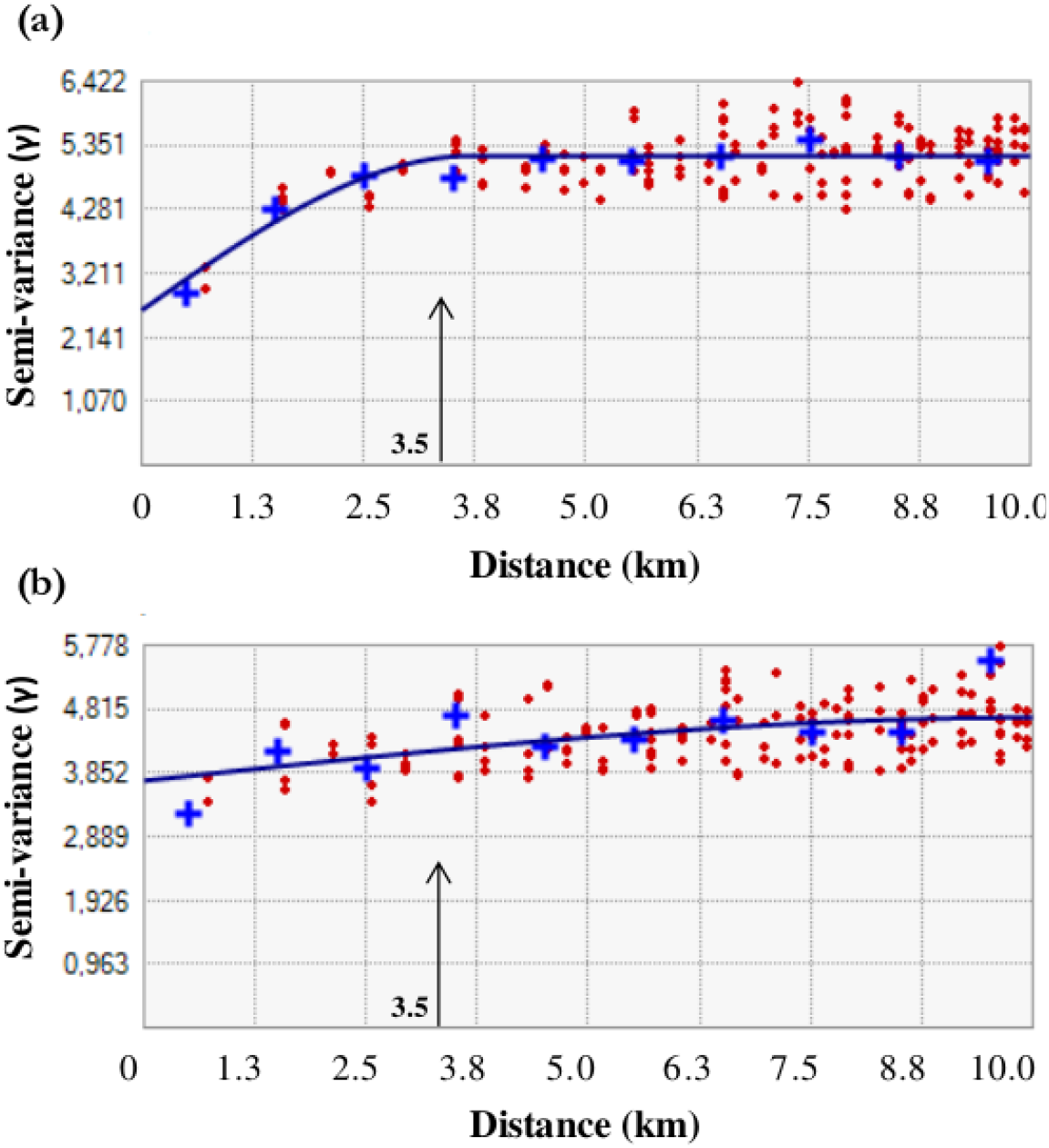
Variogram for the “reference” floristic plots in (a) coniferous and (b) braodleaved forests. The solid blue line describes a fitted spherical model. The threshold distance of 3.5 km (highlighted by arrows in the figures) was selected considering that bioindicated C:N values within this radius were spatially autocorrelated in both forest types.

**Figure S2:**
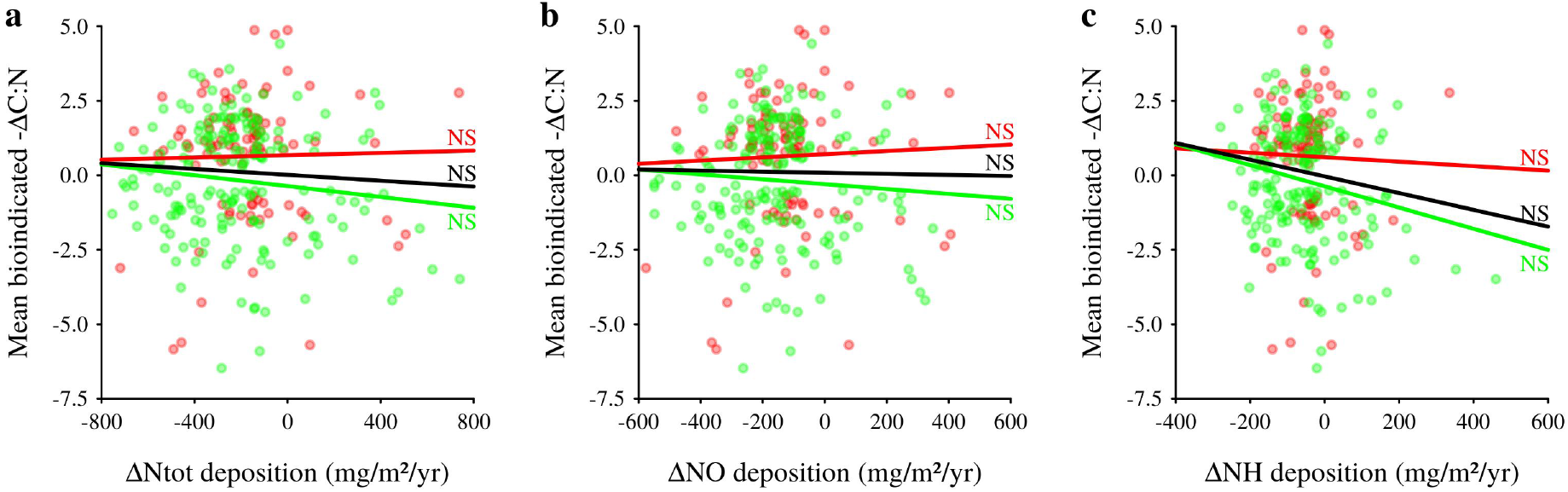
Relationships between changes in mean bioindicated C:N and total nitrogen (a), nitrogen oxyde (b) and ammonia (c) atmospheric depositions within EMEP grids. Relationships are only shown for EMEP grids with significant bioindicated ΔC:N (n = 274; see Fig. 5). Changes in bioindicated C:N and N depositions were computed as the differences between values of the reference (i.e. the 2005–2010 period) and former periods (ΔC:N = C:N_reference_ – C:N_former_; ΔNdepo = Ndepo_reference_ – Ndepo_former_). As N content decrease with C:N, we multiplied ΔC:N values by −1 in order to an increase or a decrease in bioindicated N content and deposition vary in a positive way. Nitrogen deposition values were predicted by the EMEP model (EMEP, 2011). Red and green points are observations in coniferous and broadleaved forests, respectively. Red, green and black lines are linear models fitted in coniferous, broadleaved forests, and whatever the forest type, respectively. Linear model significances were tested by comparing the slope to 0 through a Student’s *t* test (*: *P* < 0.05; NS for non significance : *P* ≥ 0.05).

**Figure S3:**
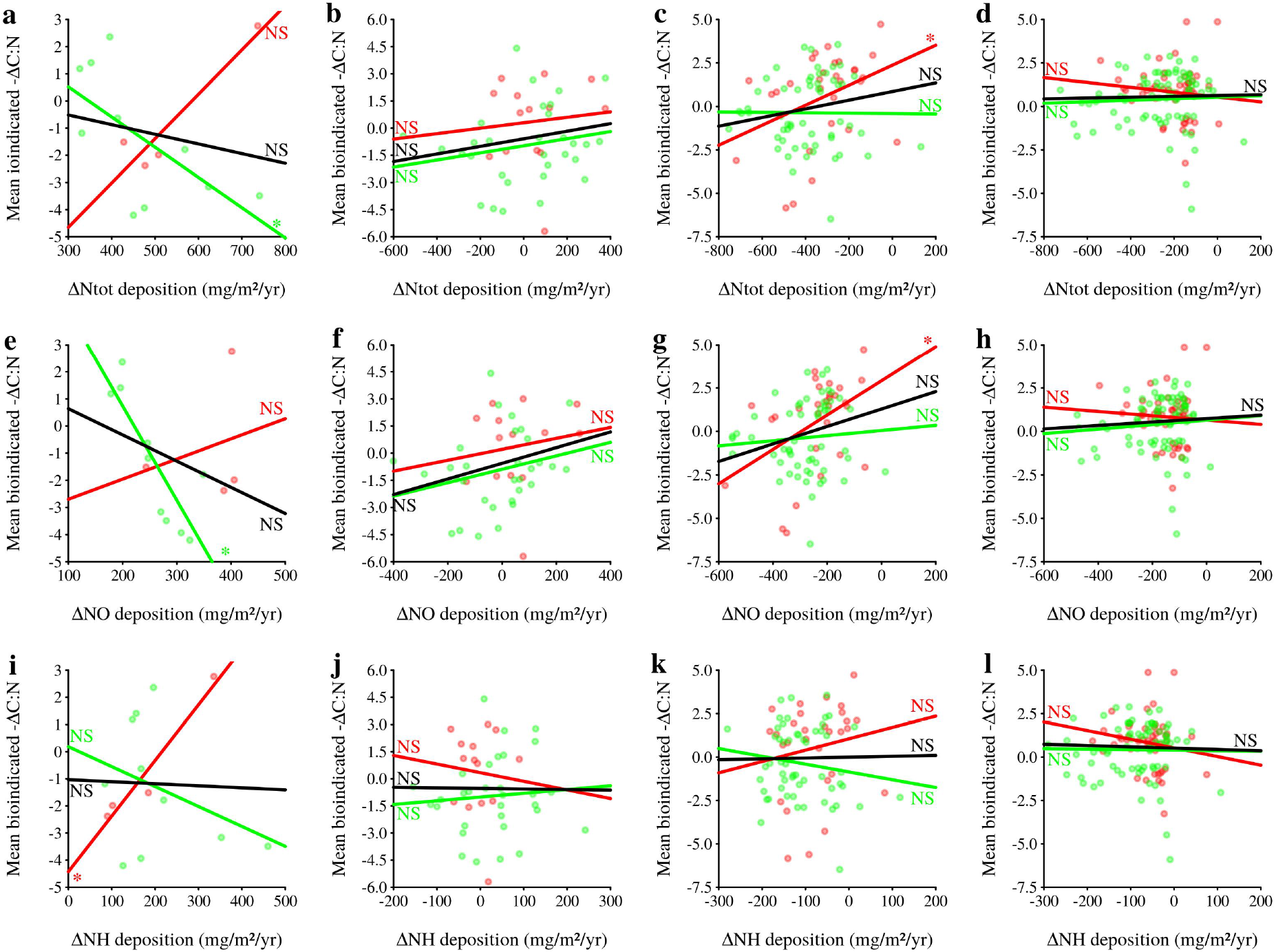
Temporal variation in relationships between changes in mean bioindicated C:N and total nitrogen (a, b, c, d), nitrogen oxyde (e, f, g, h) and ammonia (i, j, k, l) atmospheric depositions within EMEP grids. Panels a, e and i show relationships for the 1910–1949 period. Panels b, f and j show relationships for the 1950–1974 period. Panels c, g and k show relationships for the 1975–1989 period. Panels d, h and l show relationships for the 1990–2004 period. Relationships are only shown for EMEP grids with significant bioindicated ΔC:N (see Fig. 5). Changes in bioindicated C:N and N depositions were computed as the differences between values of the reference (i.e. the 2005–2010 period) and former periods (ΔC:N = C:N_reference_ – C:N_former_; ΔNdepo = Ndepo_reference_ – Ndepo_former_). As N content decrease with C:N, we multiplied ΔC:N values by −1 in order to an increase or a decrease in bioindicated N content and deposition vary in a same way. Nitrogen deposition values were predicted by the EMEP model (EMEP, 2011). Red and green points are observations in coniferous and broadleaved forests, respectively. Red, green and black lines are linear models fitted in coniferous, broadleaved forests, and whatever the forest type, respectively. Linear model significances were tested by comparing the slope to 0 through a Student’s *t* test (*: *P* < 0.05; NS for non significance : *P* ≥ 0.05).

**Figure S4:**
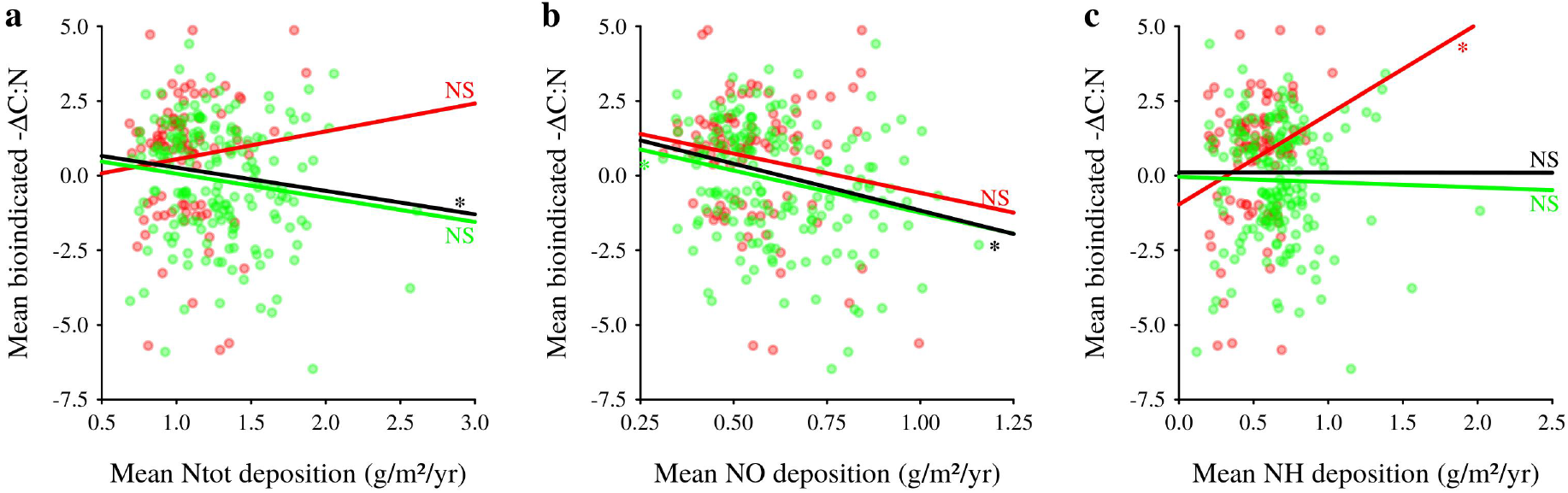
Relationships between changes in mean bioindicated C:N and mean total nitrogen (a), nitrogen oxyde (b) and ammonia (c) atmospheric depositions within EMEP grids. Relationships are only shown for EMEP grids with significant bioindicated ΔC:N (n = 274; see Fig. 5). Changes in bioindicated C:N were computed as the differences between values of the reference (i.e. the 2005–2010 period) and former periods (ΔC:N = C:N_reference_ – C:N_former_. Mean N atmospheric deposition were computed as prediction average of the EMEP model (EMEP, 2011) between the reference and the former periods. As N content decrease with C:N, we multiplied ΔC:N values by −1 in order to an increase or a decrease in bioindicated N content and in mean N deposition vary in a same way. Red and green points are observations in coniferous and broadleaved forests, respectively. Red, green and black lines are linear models fitted in coniferous, broadleaved forests, and whatever the forest type, respectively. Linear model significances were tested by comparing the slope to 0 through a Student’s *t* test (*: *P* < 0.05; NS for non significance : *P* ≥ 0.05).

**Figure S5:**
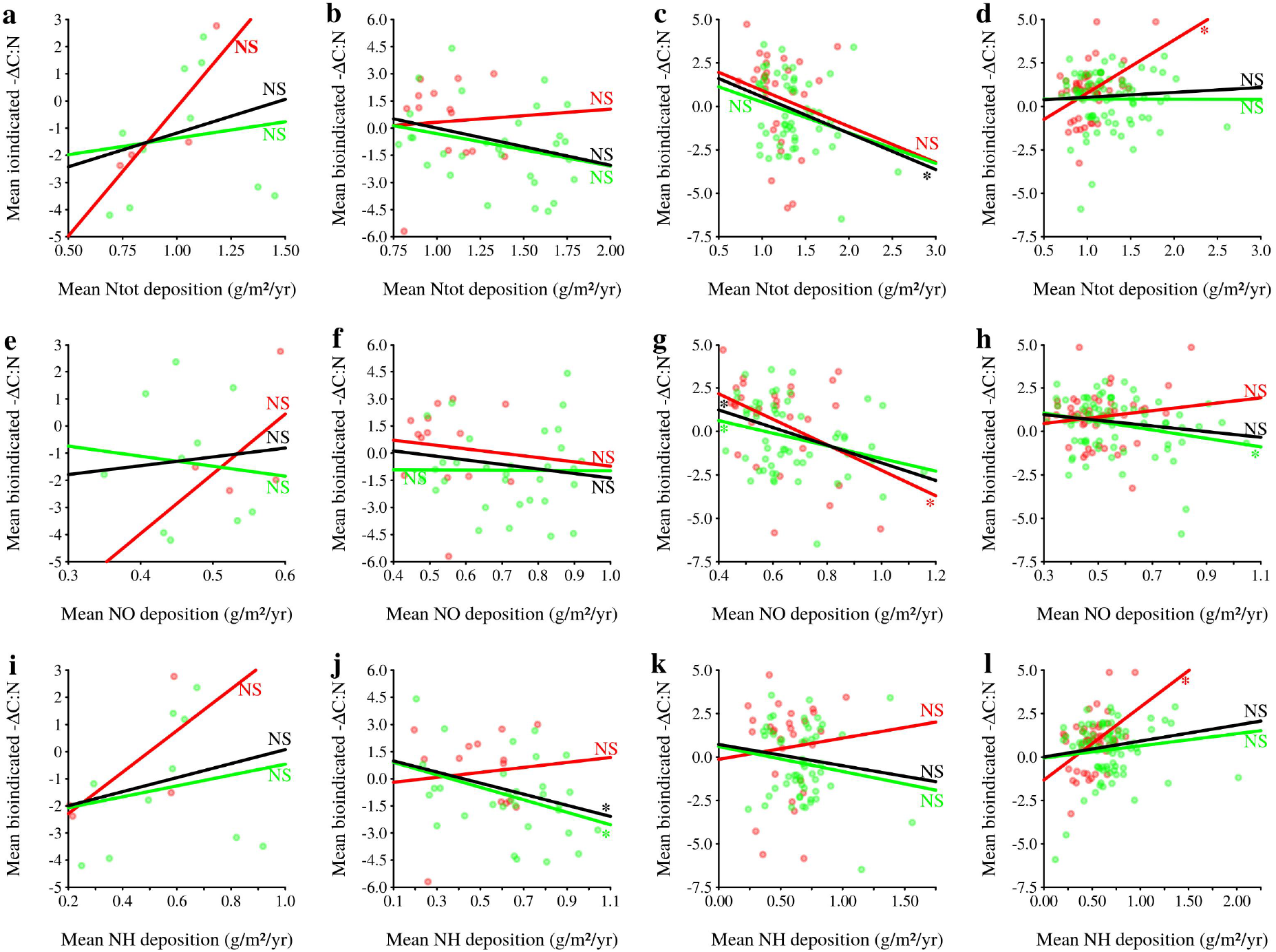
Temporal variation in relationships between changes in mean bioindicated C:N and mean total nitrogen (a, b, c, d), nitrogen oxyde (e, f, g, h) and ammonia (i, j, k, l) atmospheric depositions within EMEP grids. Panels a, e and i show relationships for the 1910–1949 period. Panels b, f and j show relationships for the 1950–1974 period. Panels c, g and k show relationships for the 1975–1989 period. Panels d, h and l show relationships for the 1990–2004 period. Relationships are only shown for EMEP grids with significant bioindicated ΔC:N (see Fig. 5). Changes in bioindicated C:N were computed as the differences between values of the reference (i.e. the 2005–2010 period) and former periods (ΔC:N = C:N_reference_ – C:N_former_. Mean N atmospheric deposition were computed as the prediction average of the EMEP model (EMEP, 2011) between the reference and the former periods. As N content decrease with C:N, we multiplied ΔC:N values by −1 in order to an increase or a decrease in bioindicated N content and in mean N deposition vary in a same way. Red and green points are observations in coniferous and broadleaved forests, respectively. Red, green and black lines are linear models fitted in coniferous, broadleaved forests, and whatever the forest type, respectively. Linear model significances were tested by comparing the slope to 0 through a Student’s *t* test (*: *P* < 0.05; NS for non significance : *P* ≥ 0.05).

**Figure S6:**
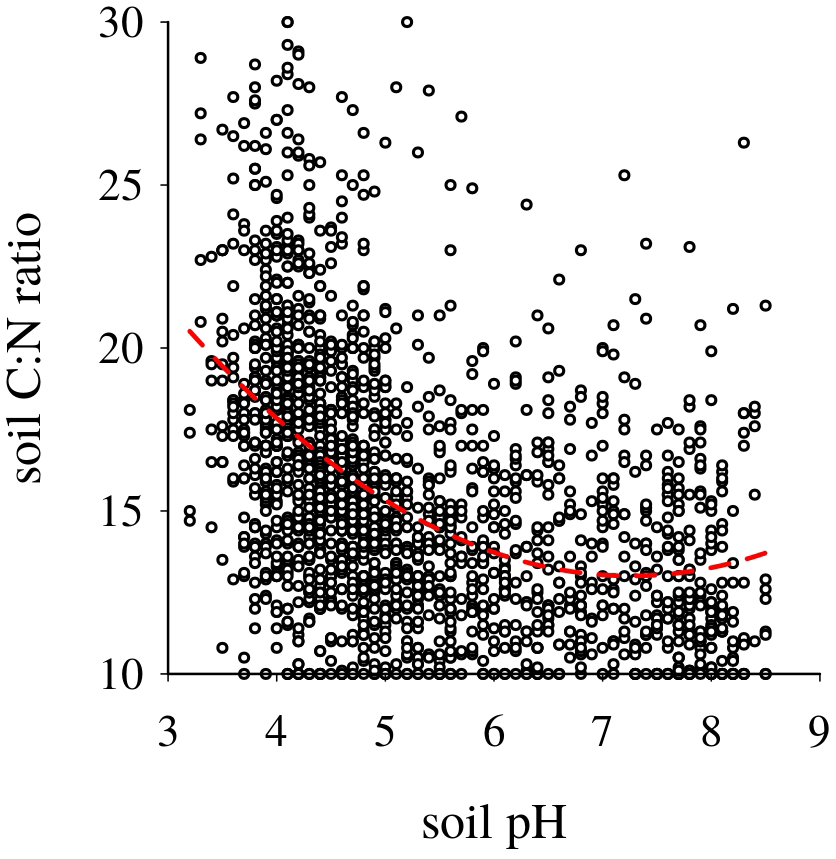
Relationship between measured soil pH and C:N ratio. The red line is the linear model fitted from 1907 observations extracted from the EcoPlant database.

